# A pseudoenzyme controls the architecture of the extreme distal tip of motile cilia

**DOI:** 10.64898/2026.02.04.703868

**Authors:** Juyeon Hong, Sidharth Ramu Nair, Eirene L. Jo, Rebecca L. Young, John B. Wallingford

## Abstract

Motile cilia are evolutionarily conserved organelles performing essential roles in development and tissue homeostasis. Unlike the core scaffold of motile cilia, the distal regions remain relatively less explored and display great diversity across species. Here, we describe a previously uncharacterized ciliary protein Jhc1, localizing at the extreme distal tip of multiciliated cell (MCC) cilia and encoded only in the genomes of non-mammalian vertebrates. Jhc1 is essential for cilia structure and length in *Xenopus*, an activity that is conserved in Jhc1 from reptiles and fish. Phylogenetic analysis and structure modeling suggests that Jhc1 arose by duplication and neofunctionalization of thiamine triphosphatase, and we show that residues crucial for that enzyme’s function have been lost and replaced by new residues essential for ciliary localization. These data provide new insights into the molecular mechanisms underlying the broad diversification of the structures at the tip of motile cilia during evolution.

## Introduction

Cilia are evolutionarily conserved organelles present in cells from unicellular algae to humans^1,2^. In mammals, motile cilia serve essential roles in generating fluid flow in the central nervous system, clearing mucus in the respiratory tract, and transporting gametes in the reproductive tracts^1^. In other animals, motile cilia play a wide range of additional functions. In amphibians, for example, cilia-mediated fluid flow promotes gas exchange on the epidermis also prevent microorganisms and debris from attaching to the skin^3^. In fish, motile cilia aid in odor detection by promoting dynamic fluid exchange between the environment and the nose^4^. Corals use motile cilia to transport oxygen from the surface deeper into the reef^5^.

How motile cilia are specialized for such clade-specific functions remains poorly understood. Indeed, the core structure of motile cilia is thought to have changed little since the last eukaryotic common ancestor. For example, the structure of the microtubule-based axoneme and basal body are well-conserved^6^. So, too, are molecular machines that build and maintain cilia including intraflagellar transport (IFT) system, dynein motors, and docking complexes^7,8^. In fact, some lineages that lack cilia today, such as plants or fungi, still retain remnants of ciliary genes, suggesting a secondary loss of cilia during their evolution^9,10^.

Unlike these core machineries, the distal ends of cilia display intriguing diversity across different species, cell types, and even developmental stages^11^. In unicellular organisms such as *Chlamydomonas* and *Tetrahymena*, the central pair microtubules extend beyond the microtubule doublets and are inserted into a cap which is attached to the end of the ciliary membrane^12-15^. In mammalian epithelia lining the trachea or oviduct, an amorphous tip caps the central pair as well as A-tubule singlets linking these to a ciliary crown constituted by fibrils emanating from the ciliary membrane^16-18^. The ciliary tip in adult frog palate shows two asymmetrical plates^19^, while the tip of motile cilia in tadpole epidermis consists of a bulb-like structure^20^. Little else is known of the structure or assembly of the EDT in any species, and nothing is known of molecular mechanisms that mediate species-specific specializations of the distal tips of motile cilia.

One mechanism for the evolution of such novelties is the duplication and neofunctionalization of genes^21,22^, and evidence suggests this is the case for cilia, as several key ciliary proteins are non-catalytic pseudoenzymes. For example, Ulk4 (Unc-51-like kinase) progressively lost essential catalytic residues during evolution^23^ but still plays a critical role in motile ciliogenesis. Loss of ULK4 causes ciliopathy-related phenotypes including hydrocephalus and brain malformation^24-26^. Likewise, the pseudophosphatase Styxl1 is required to scaffold chaperones during sperm axoneme assembly^27^, and several members of the NME family of nucleoside kinases have lost catalytic activity during evolution and play structural roles in cilia^28^.

Here, we identified the uncharacterized protein Jhc1 as a component of the EDT of *Xenopus* motile cilia. Notably, Jhc1 is not present in mammals but is essential for normal control of cilia tip structure and cilia length in frogs. Its function is likely conserved, as homologues from fish and reptiles can rescue the effect of Jhc1 loss. Sequence comparison and structural modelling revealed that Jhc1 shares a similar architecture with thiamine triphosphatase (Thtpa) but key catalytic residues of Thtpa are absent in Jhc1, and at least one of these residues is essential for Jhc1 localization to cilia. Our data suggest that Jhc1 arose from an ancient duplication of Thtpa followed by functional divergence and represents a lineage-specific novelty that shapes the distal ciliary architecture of motile cilia in non-mammalian vertebrates.

## Results

### Jhc1 localizes to the extreme distal tip of motile cilia in *Xenopus* MCCs

We previously identified a spatially distinct domain at the ends 9+2 cilia on multiciliated cells (MCCs) of *Xenopus* mice, and humans that is labelled by two proteins, Ccdc78 and Ccdc33.^29^ We refer to this domain as the extreme distal tip (EDT) of motile cilia because in *Xenopus*, these proteins specifically mark a region of ∼0.2μm that caps the larger, well-characterized domain marked by the central pair protein Spef1 (Fig. 1A)^29-31^.

**Figure 1.**
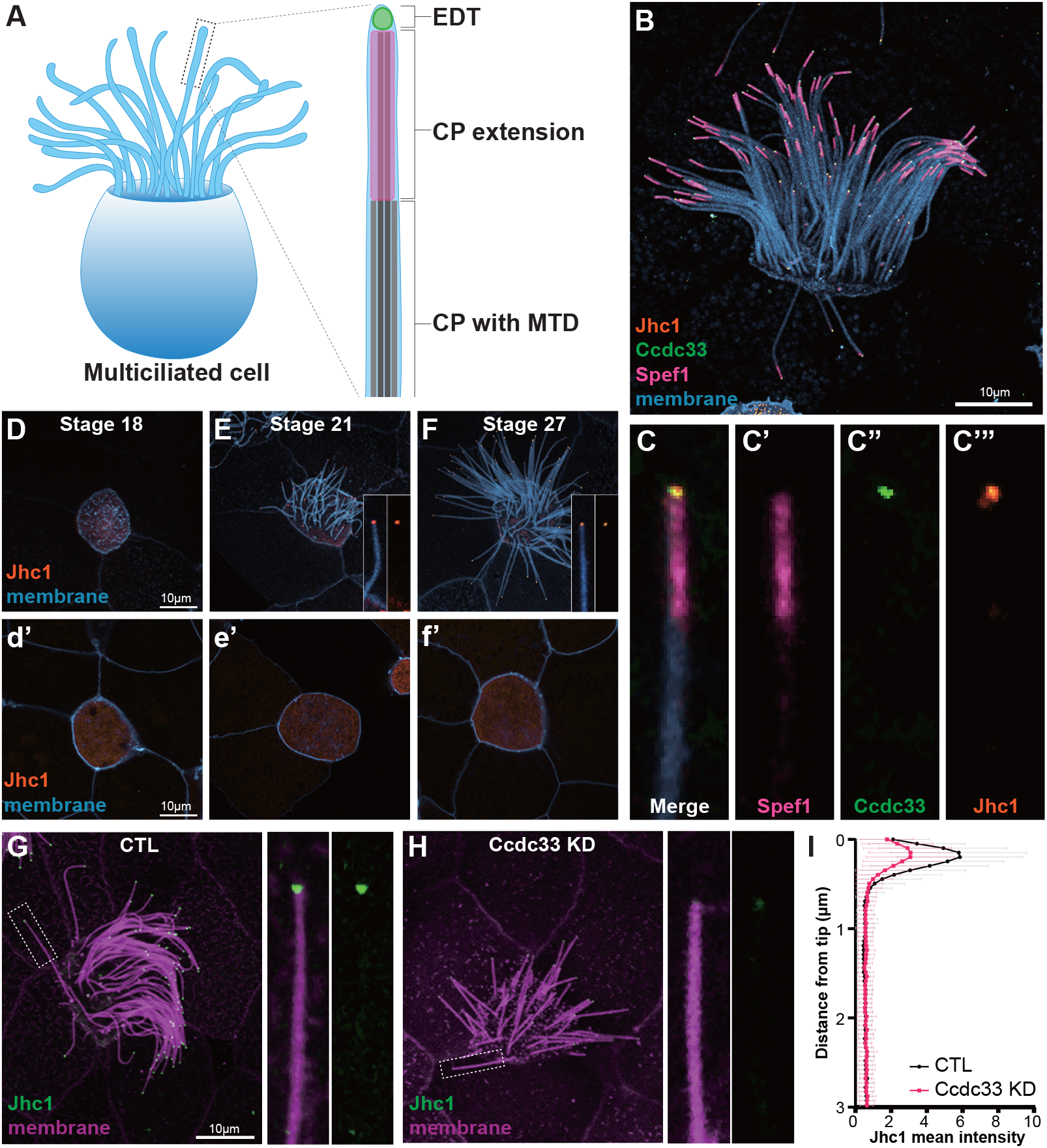
Jhc1 localizes to the extreme distal tip of motile cilia in *Xenopus* MCCs. (A) Schematic image of MCC with the magnified view of distal ciliary region, marked with extreme distal tip (EDT), Central pair extension (CP extension) and Central pair (CP) with microtubule doublets (MTD). (B) *Xenopus* MCC expressed with membrane-BFP (blue), Spef1-RFP (magenta), GFP-Ccdc33 (green) and Halo-Jhc1 (orange). Scale bar represents 10μm. (C) Magnified view of a cilium from figure 1B. (D-F) Image of *Xenopus* MCC cilia and cytosol expressed with membrane-BFP (blue) and GFP-Jhc1 (orange) during MCC development; before cilia protrusion (stage 18, D), during ciliogenesis (stage 21, E), mature cilia (stage 27, F). The magnified view of the cilium is inserted at the bottom right. Scale bar represents 10μm. (G-H) Image of *Xenopus* MCC expressed with GFP-Jhc1 (green) and membrane-RFP (magenta) in control (G) and Ccdc33 KD (H) embryos. The magnified view of the cilium is shown on the right. Scale bar represents 10μm. (I) Quantification of GFP-Jhc1 intensity along the axoneme of control and Ccdc33 KD MCCs, normalized by average intensity. Graph shows mean ± SD (n=120 cilia, 40 cells, 10 embryos, 3 experiments).

In proteomic experiments using *Xenopus* MCCs, we found that Ccdc33 interacts with the uncharacterized protein Loc108698169^20^. Based on the data below, we propose to rename this protein ‘Just the Head of Cilia 1’ (Jhc1). We expressed Halo-tagged Jhc1 in *Xenopus* MCCs and found it colocalized with GFP-Ccdc33 at the EDT, just distal to Spef1-RFP (Fig. 1B-C)^13,30-32^. As observed for other EDT proteins^20^, Halo-Jhc1 showed consistent localization at the EDT in both developing/elongating cilia and in full-length homeostatic cilia (Fig. 1D-F).

To explore the functional relationships in the EDT, we knockdown (KD) Ccdc33 using a previously validated morpholino-oligonucleotide^29^. We found that Ccdc33 KD significantly decreased intensity of GFP-Jhc1 at the tip of MCC cilia as compared to the controls (Fig. 1G-I). These data suggest that Jhc1 is a novel component of the EDT, so we explored the evolution of this wholly uncharacterized protein.

### Jhc1 is conserved across non-mammalian vertebrates and shares structural similarity to the thiamine triphosphatase, Thtpa

Jhc1 was not annotated in the *Xenopus* model organism database, Xenbase^33^, and all homologues identified with BLASTp were likewise uncharacterized (Supp. Table. 1). Interestingly, Jhc1-like proteins were present in diverse vertebrates, including amphibians, reptiles, and both cartilaginous and bony fish, but were absent from mammals and non-vertebrate animals (Fig. 2A, Supp. Table. 1). To ask if the proteins identified by BLASTp represent true orthologues of Jhc1, we examined the genomic synteny of the encoding genes^34^. *Xenopus jhc1* is located within the first intron of another gene, *ralgps1*.*L* on chromosome 8; it lies adjacent to *angptl2*.*L* (Supp. Fig. 1A). Notably, this genomic location for *jhc1* was precisely conserved in several other vertebrates, including fish and two reptiles (Supp. Fig. 1B-D). By contrast, no *Jhc1* gene is present in this syntenic region in humans and mice (Supp. Fig. 1E-F).

**Figure 2.**
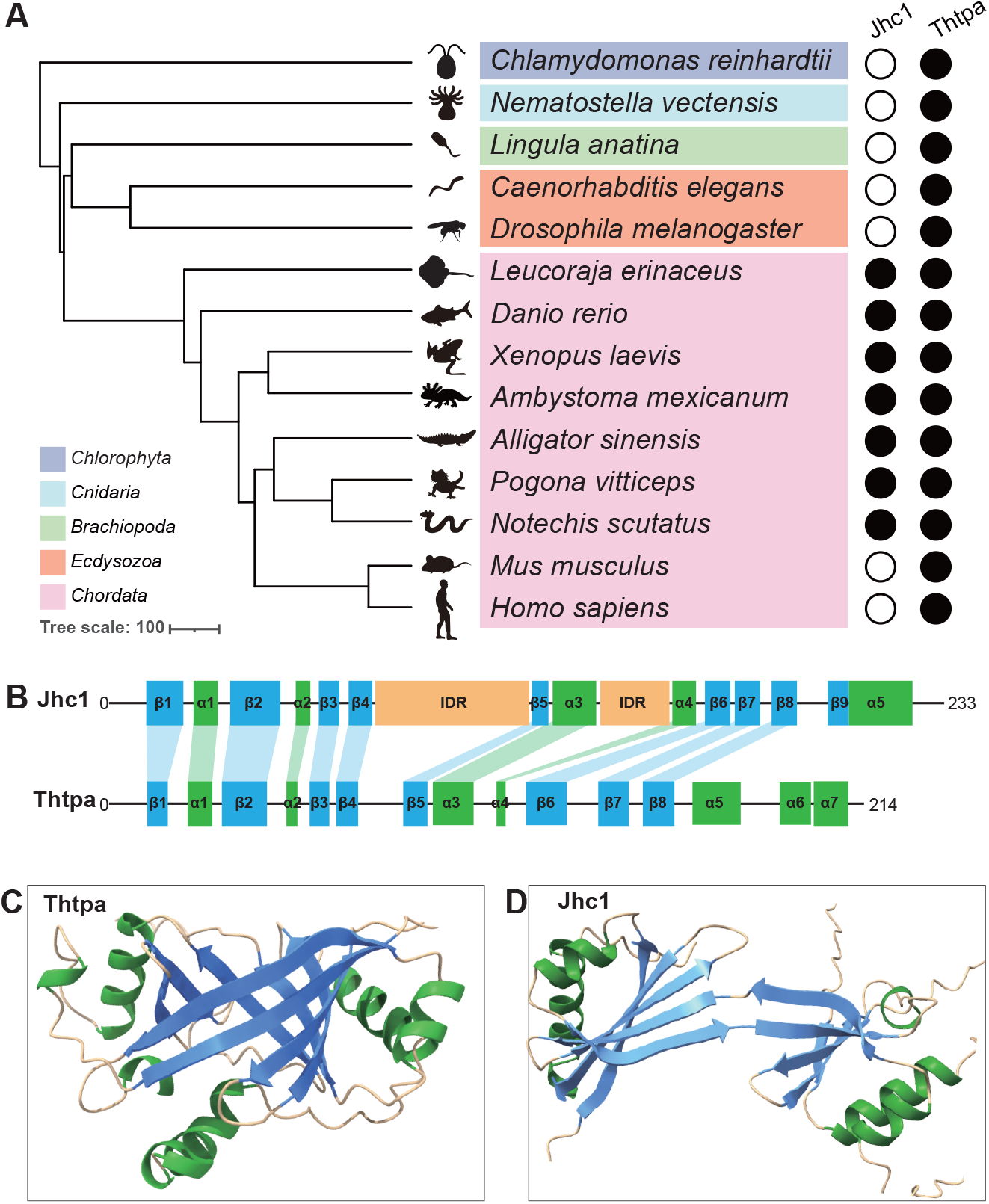
Jhc1 displays a similar structure to Thtpa. (A) The presence (filled circle) of Jhc1 and Thtpa indicated in diverse species. (B) Alignment of secondary structure of Jhc1 and Thtpa. α-helix is colored in green, β-sheet in blue and IDR in orange. (C and D) Predicted protein structure of Jhc1 (D) and Thtpa (C) from Alphafold3. The IDR of Jhc1 (top right) is hidden. α-helix is colored in green, β-sheet in blue and IDR in orange.

To identify proteins related to Jhc1, we performed PSI BLAST against NCBI Protein reference sequences^40^ from species across the animal tree of life (see Methods). This analysis identified additional Jhc1 orthologues and, with lower scores, also identified thiamine triphosphatases (Thtpa) from several species (Supp. Table. 2). Phylogenetic analysis of the output of this search placed Jhc1 in a strong monophyletic clade, clearly distinct from Thtpa (Supp Fig. 2A). Specifically, Jhc1 sequences clustered together across vertebrates with high confidence, whereas Thtpa sequences formed a weakly resolved assemblage with interspersed relationships (Supp. Fig. 2A). A member of the CYTH-superfamily, Thtpa is an enzyme that breaks down thiamine triphosphate into thiamine diphosphate and is widely conserved across eukaryotes (Fig. 2A)^36-38^. Synteny confirmed that Thtpa is encoded by an independent gene at a conserved genomic location that is distinct from *jhc1* (Supp. Fig. 3A-E).

Jhc1 and Thtpa share highly similar secondary structures, with β-sheets and α-helixes dispersed along their length, though Jhc1 is distinguished by two centrally located intrinsically disordered regions (Fig. 2B). This secondary structure is highly conserved in both proteins across species (Supp. Fig. 4). The tertiary structure of human Thtpa has been solved, revealing that like other CYTH domain proteins the repeated sheets form a β-barrel (PDB: 3TVL)^38^. That structure is precisely reflected in an Alphafold3^39^ prediction of *Xenopus* Thtpa (Fig. 2C, Supp. Fig. 5), while by contrast, *Xenopus* Jhc1 is confidently predicted to adopt an open conformation (Fig. 2D). Jhc1 from fish and alligator are predicted to adopt similar structures (Supp. Fig. 6A and 6B).

These distinct structures led us to a final test of the relationship between Jhc1 and CYTH domain proteins such as Thtpa. Analysis with HHpred identified CYTH-family proteins such as the triphosphate tunnel metalloenzyme (PBD: 7NS9_A) as the strongest remote homologues of Jhc1 in distantly related species (Supp. Table. 3).

### Jhc1 is expressed specifically in multiciliated tissues

To learn more about *jhc1*, we next examined its expression and compared it to *thtpa* in *Xenopus* using whole mount in situ hybridization. *jhc1* was expressed in two different populations of MCCS, those on the epidermis and those in the nephrostomes of the pronephros (Fig. 3A). In the epidermis, specific expression in MCCs was confirmed by co-labeling with acetylated tubulin (Fig. 3A, C). By contrast, *thtpa* was not enriched in MCCs but displayed weak, broad expression throughout the embryo (Fig. 3B).

**Figure 3.**
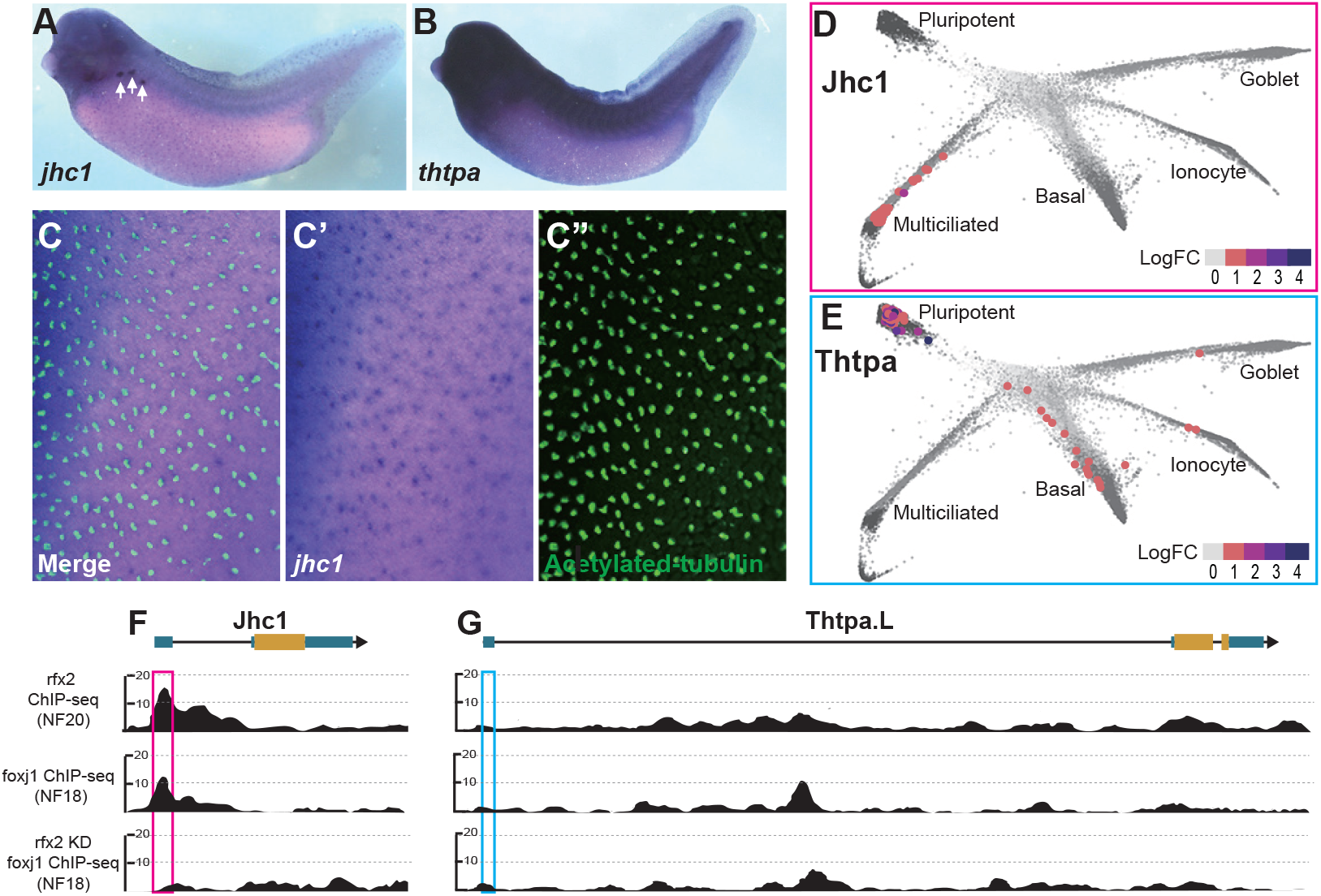
Jhc1 and Thtpa display distinct expression patterns. (A and B) Whole mount in situ hybridization (WISH) of Jhc1 (A) and Thtpa (B) in *Xenopus* embryo. White arrows on Jhc1 WISH embryo indicate pronephros tubule. (C) Colocalization of acetylated-tubulin immunostaining and Jhc1 WISH signal in *Xenopus* embryo MCC. (D and E) tSNE plot of *Xenopus* animal cap scRNA-seq captured from UCSC cell browser with *Xenopus* animal cap data (https://cells-test.gi.ucsc.edu/?ds=xenopus-dev&gene=gene16200#). Jhc1 expression is shown in multiciliated cells (D). Thtpa expression is shown in pluripotent, epithelial progenitors, basal cells, ionocyte and goblet cells (E). Data are present with log fold change. (F and G) Browser screenshot of Jhc1 (F) and Thtpa (G) genes with ChIP-seq profiles from Xenbase JBrowse (https://xenbase.org/xenbase/displayJBrowse.do?data=data/xl10_1). Top track shows Rfx2 binding profile, middle track shows Foxj1 binding profile, bottom track shows Foxj1 binding profile in Rfx2 knockdown embryos.

To confirm these results, we examined single-cell RNA-seq data from *Xenopus* embryo animal caps^41^. We found Jhc1 to be strongly expressed only in the MCC lineage (Fig. 3D), whereas Thtpa was broadly detected throughout the tissue, and was enriched in pluripotent progenitors and basal cells, but not in MCCs (Fig. 3E).

Finally, we examined *Xenopus* ChIP-seq profiles^42,43^, finding that the promoter region of *jhc1* is bound strongly by Rfx2 and Foxj1, key transcription factors that drive MCC differentiation (Fig. 3F)^42-44^. Moreover, Foxj1 binding was lost from the *jhc1* promoter after knockdown of Rfx2 (Fig. 3F), consistent with *jhc1* being under control of the conserved MCC transcriptional circuit^43^. By contrast, the promoter region of *thtpa* was not bound by these factors (Fig. 3G). Together, the divergent structures and gene expression patterns suggest that *jhc1* performs a specific function in MCCs that is not shared with the related gene, *thpta*. We next tested this idea directly.

### Jhc1 is required for distal ciliary structure and cilia length of MCC cilia

The EDT proteins Ccdc78 and Ccdc33 are required to regulate the molecular organization of the distal region of MCC cilia and as well as cilia length^13,29^. To ask if Jhc1 performs a similar function, we performed Jhc1 knockdown. As observed for Ccdc33^29^, loss of Jhc1 resulted in a reduction in cilia length (Fig. 4A, B, E) as well as a partial loss of the Spef1-labeled central pair extension (Fig. 4a′, 3b′ and F). Importantly, loss of the Spef1 domain cannot account for the reduced cilia length, as overall length is reduced by over 5 microns, yet the Spef1 domain is reduced only by ∼1 micron (Fig. 4E, F). Crucially, the defects in both cilia length and Spef1 localization were rescued by re-expression of *Xenopus* Jhc1, confirming the specificity of morpholino (Fig. 4C, E, F). By contrast, expression of Thtpa did not rescue the Jhc1 knockdown effects on cilia length or Spef1 localization (Fig. 4D, E, F), consistent with the idea that the molecular function of Jhc1 is not shared with Thtpa.

**Figure 4.**
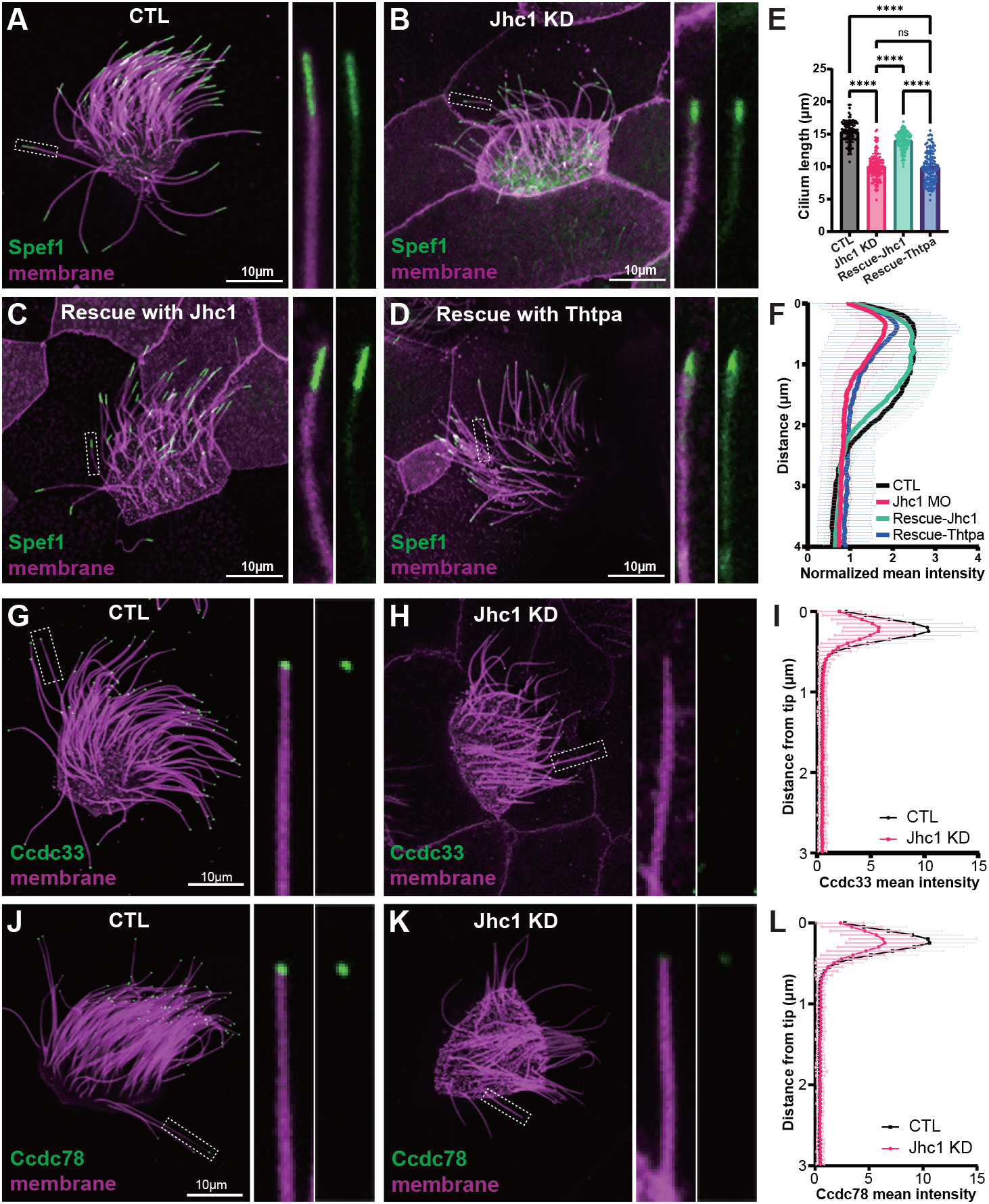
Jhc1 is required for distal ciliary structure and length of MCC cilia. (A-D) Image of *Xenopus* MCC expressed with membrane-RFP (magenta) and Spef1-GFP (green) in control (A), Jhc1 KD (B), rescued with Jhc1 (C) and rescued with Thtpa (D). The magnified view of the cilium is shown on right. Scale bar represents 10μm. (E) Quantification of cilia length in control, Jhc1 KD, rescued with Jhc1 and rescued with Thtpa *Xenopus* MCC. Graph shows mean ± SD and analyzed with Ordinary one-way ANOVA multiple comparisons (n=119 cilia, 40 cells, 10 embryos, 3 experiments). P****<0.0001. (F) Quantification of Spef1-GFP fluorescent intensity along the axoneme of control, Jhc1 KD, rescued with Jhc1 and rescued with Thtpa. Graph shows mean ± SD (n=119 cilia, 40 cells, 10 embryos, 3 experiments). (G-H) Image of *Xenopus* MCC expressed with GFP-Ccdc33 (green) and membrane-RFP (magenta) in control (J) and Jhc1 KD (K) embryos. The magnified view of the cilium is shown on the right. Scale bar represents 10μm. (I) Quantification of GFP-Ccdc33 intensity along the axoneme of control and Jhc1 KD MCCs, normalized by average intensity Graph shows mean ± SD (n=120 cilia, 40 cells, 10 embryos, 3 experiments). (J-K) Image of *Xenopus* MCC expressed with GFP-Ccdc78 (green) and membrane-RFP (magenta) in control (J) and Jhc1 KD (K) embryos. The magnified view of the cilium is shown on the right. Scale bar represents 10μm. (L) Quantification of GFP-Ccdc78 intensity along the axoneme of control and Jhc1 KD MCCs, normalized by average intensity Graph shows mean ± SD (n=120 cilia, 40 cells, 10 embryos, 3 experiments).

We next explored the effect of Jhc1 loss on the EDT. In control MCC cilia, GFP-Ccdc33 and Ccdc78 mark the EDT within ∼0.2μm enrichment at the distal tip. Loss of Jhc1 reduced the intensity of both Ccdc33 and Ccdc78 at the EDT by roughly half (Fig. 1G-I). These data suggest that Jhc1 is a novel component of the EDT that interacts functionally with Ccdc33 and Ccdc78 to control the structure of the distal cilia and cilia length in MCCs. We next explored the evolution of this uncharacterized but important protein.

### Ciliary localization and function are conserved in fish and reptile Jhc1

We asked first if Jhc1 function is conserved in other non-mammalian vertebrates. To this end, we expressed fluorescently tagged Jhc1 from fish or alligator in *Xenopus* MCCs and found that each localized robustly to the EDT, similar to *Xenopus* Jhc1 (Fig. 5A, B). By contrast, Thtpa from both *Xenopus* and fish was not enriched in the EDT, though it was present diffusely along the length of the axoneme (Supp. Fig. 7).

**Figure 5.**
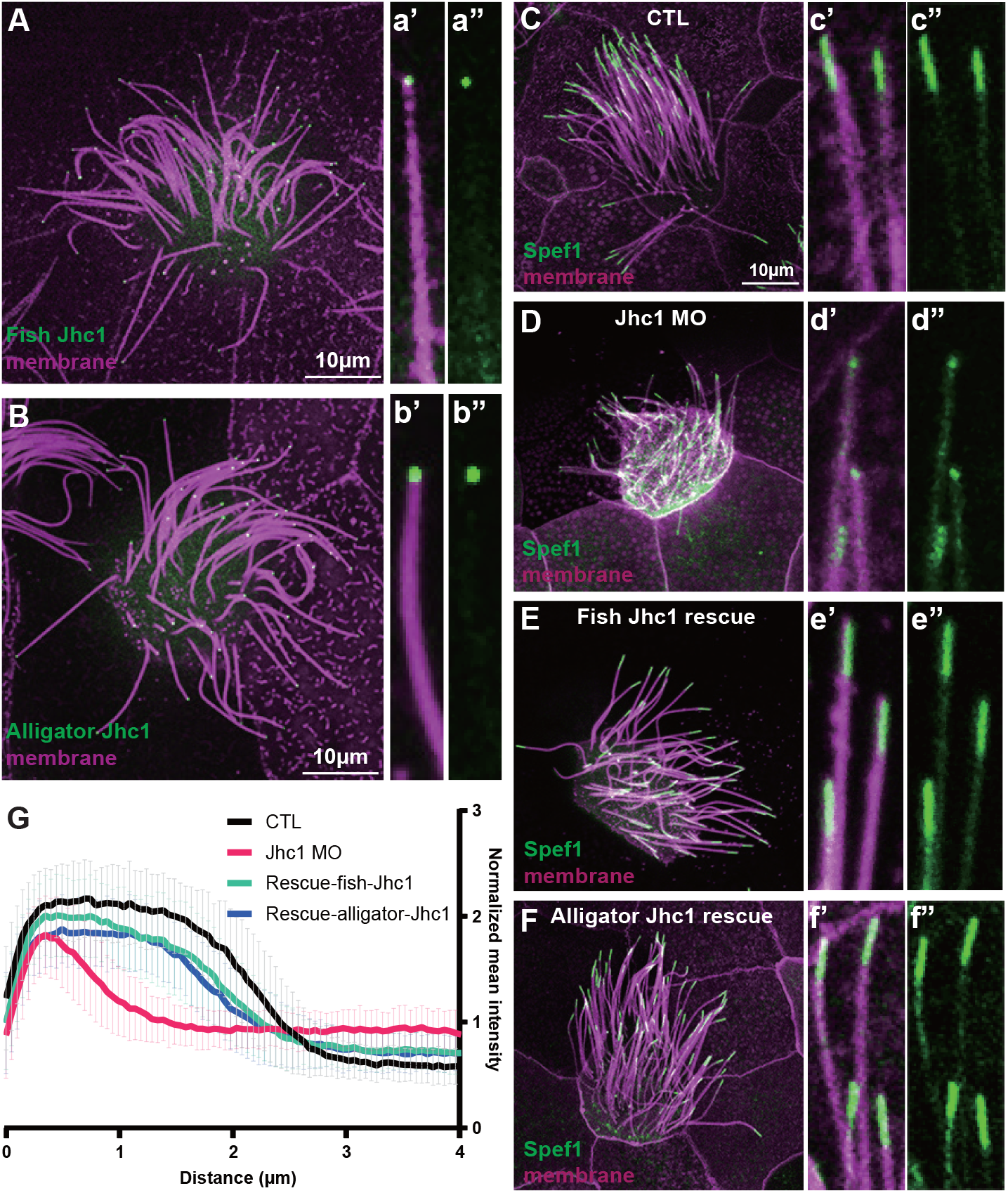
Jhc1 function is evolutionarily conserved in non-mammalian vertebrates. (A and B) *Xenopus* MCC expressed with membrane-RFP (magenta) and GFP-tagged *Danio rerio* Jhc1 (green)(A) or *Alligator sinensis* Jhc1 (green)(B). The magnified view of the cilium is shown on right. Scale bar represents 10μm. (C-F) Image of Xenopus MCC expressed with membrane-RFP (magenta) and Spef1-GFP (green) in control (C), Jhc1 KD (D), rescued with *Danio rerio* Jhc1 (E), rescued with *Alligator sinensis* Jhc1 (F). The magnified view of the cilium is shown on right. Scale bar represents 10μm. (G) Quantification of Spef1-GFP fluorescent intensity along the axoneme of control, Jhc1 KD, rescued with *Danio rerio* Jhc1 and rescued with *Alligator sinensis* Jhc1. Graph shows mean ± SD (n=120 cilia, 40 cells, 10 embryos, 3 experiments).

As more direct test of the conservation of Jhc1 function, we asked if fish or alligator Jhc1 could rescue the effect of Jhc1 loss in *Xenopus* MCCs. Indeed, they could; each effectively rescued both cilia length and Spef1 localization (Fig. 5E, F, G). These results suggest that the role for Jhc1 in MCC structure and length control is evolutionarily conserved in frogs, reptiles, and fish.

### Catalytic residues of Thtpa are missing in Jhc1

Our data suggest that Jhc1 represents a distinct gene lineage that may have evolved from Thtpa early in vertebrate evolution and taken on a ciliary function. To explore this possibility, we took advantage of detailed knowledge of the catalytic mechanism of Thtpa and other CYTH-domain proteins^36-38^. These proteins require a divalent metal cation, typically Mg^2+^, that is coordinated within the β-barrel by an ExExK motif in the first beta-sheet (Fig. 6A, B)^36,38,45^. This ExExK motif is well conserved in Thtpa across species (Fig. 5A, black box) and is confidently predicted to bind Mg^2+^ in an AlphaFold3 model of *Xenopus* Thtpa (Fig. 6B, Supp. Fig. 8A). However, the entire motif is absent in Jhc1, despite the continued presence of a β-sheet at this position (Fig. 6A, black box). When Jhc1 was modeled with Mg^2+^ using Alphafold3, the substituted residues ARG12 and GLN14 failed to form interactions, as expected (Fig. 6C, Supp. Fig. 8B).

**Figure 6.**
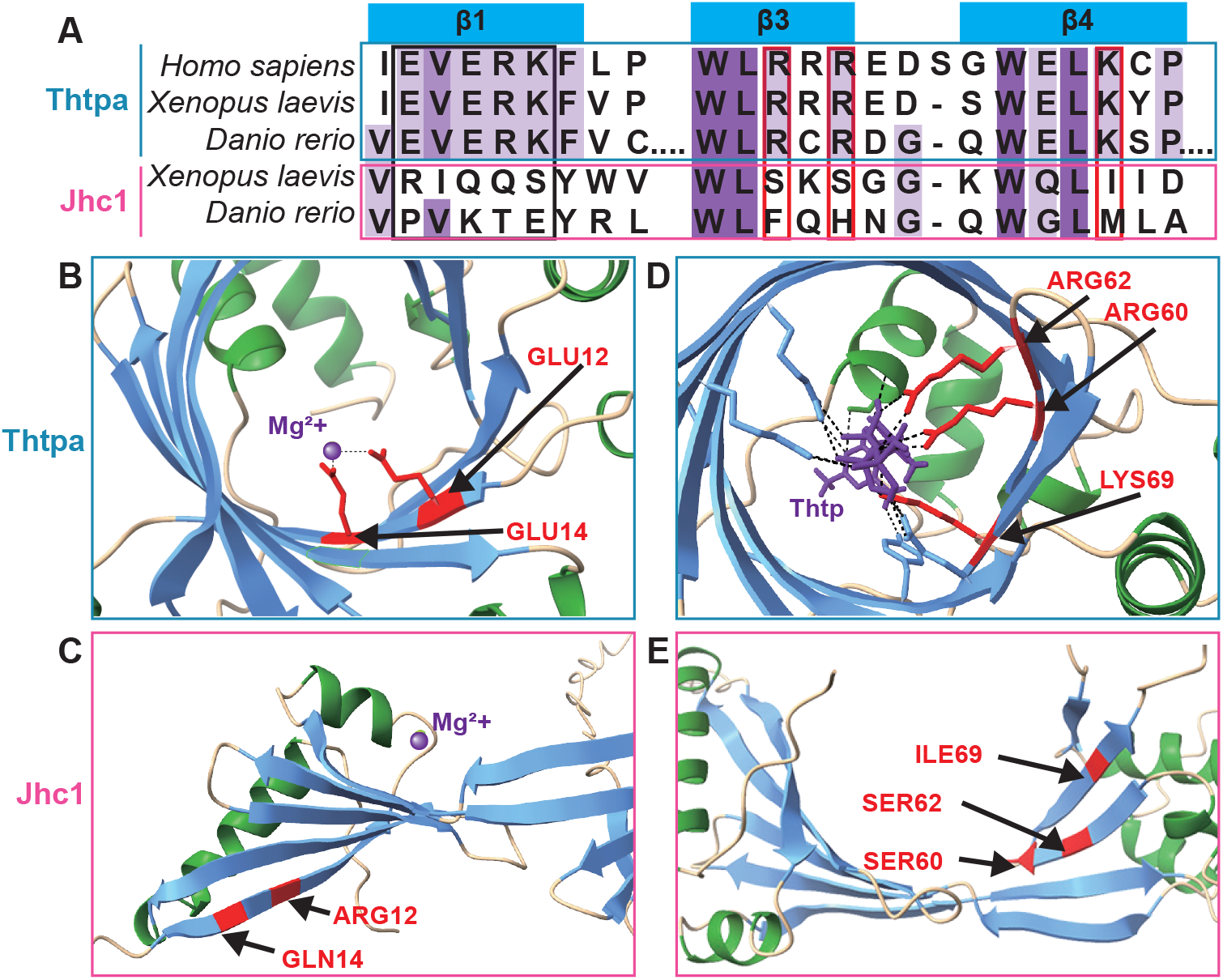
Catalytic residues of Thtpa are absent in Jhc1. (A) Amino acid sequence alignment of Thtpa (blue) and Jhc1 (magenta) from *Human, Xenopus laevis* and *Danio rerio*. Sequence similarity is shown in purple, with darker shades indicating higher similarity. Catalytic residues of Thtpa are marked with red boxes for comparison with Jhc1. (B and C) Binding of Mg2+ (purple) in Thtpa (B) and Jhc1 (C). Main interactions are labeled and highlighted in red. (D and E) Binding of Thtp (purple) in Thtpa (D) and Jhc1 (E). Main interactions are labeled and highlighted in red.

As a more stringent test, we examined basic residues essential for catalytic activity of Thtpa. Lysine-69, Arginine-55 and Arginine-57 are well conserved from fish to humans (Fig. 6A), and their mutation abrogates Thtpa activity^38^. The structure of human Thtpa (PDB: 3TVL) revealed that these basic residues coordinate interaction with triphosphate inside the β-barrel^38^. AlphaFold3 predicted precisely the same organization for *Xenopus* Thtpa (Fig. 6D, Supp. Fig. 8C, D).

Strikingly, all of these key basic residues are absent in the third and fourth β-sheets of Jhc1 in both frogs and fish (Fig. 6A), thus eliminating the possibility of catalytic activity. Moreover, AlphaFold3 revealed these residues map near one another on the same face of the open structure of *Xenopus* Jhc1 (Fig. 6E), suggesting the possibility that they mediate a novel protein-protein interaction.

### A position essential for catalysis in Thtpa performs a cilia localization function in Jhc1

Our data suggest that Jhc1 may have evolved by duplication of Thtpa followed by neofunctionalization. If this is the case, we might expect that residues no longer constrained by catalytic activity may have evolved new function related to Jhc1’s action in cilia. To explore this idea, we noted that Jhc1 interacted with Ccdc33 in proteomic experiments^29^ and that AlphaFold3 suggests that interaction is direct (Fig. 7A, B, black box). Using computational alanine scanning^46^, we identified residues critical for this interaction (Fig. 7C, D, Black box; Supp. Table. 4). Among the high-scoring residues was Isoleucine-69, which in Thtpa is the catalytic residue Lysine-69 (Fig. 6D, E and 7E).

**Figure 7.**
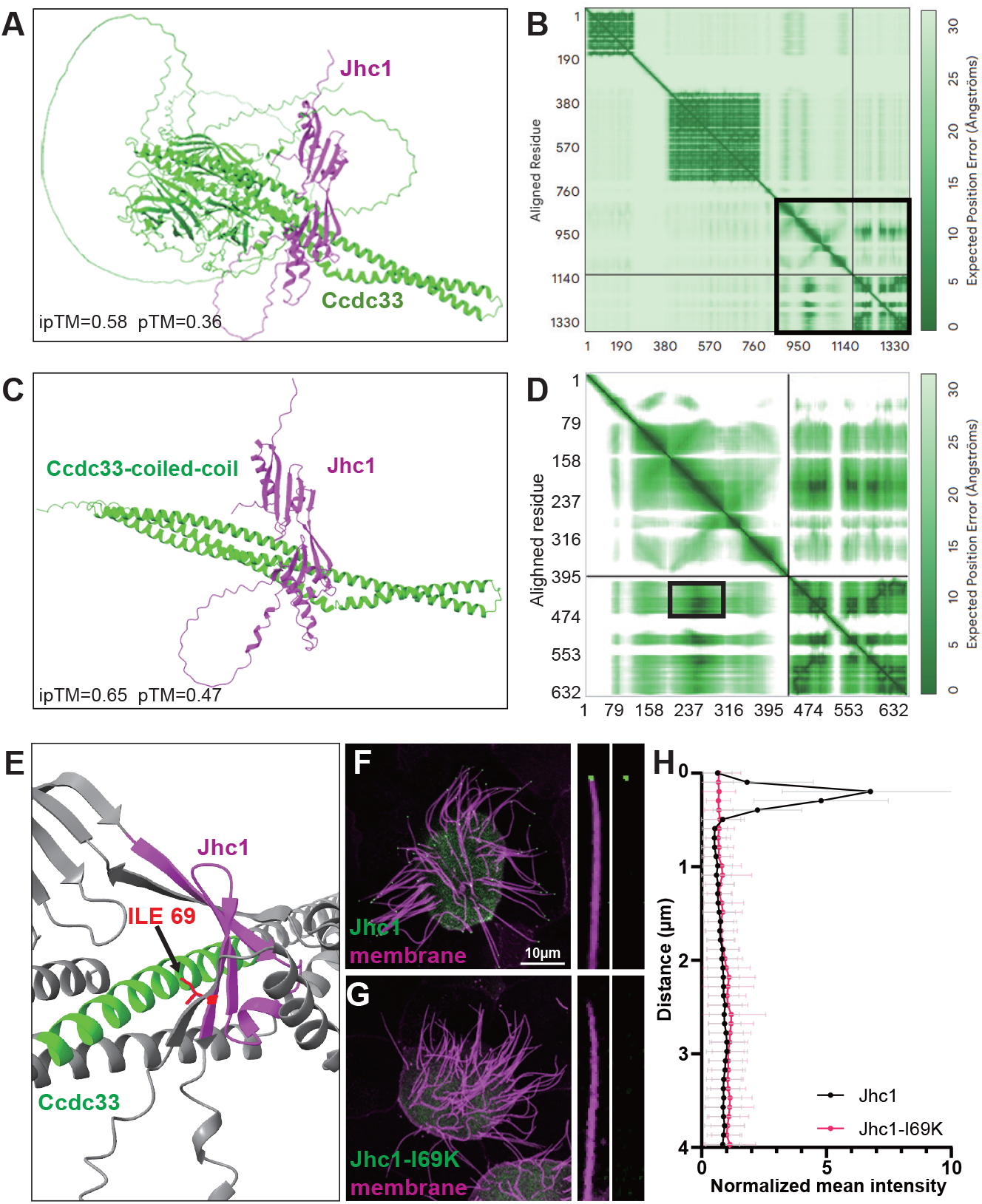
A residue essential for catalysis in Thtpa performs a cilia localization function in Jhc1. (A) Alphafold3 interaction of Jhc1 (magenta) and Ccdc33 (green). (B) PAE plot of figure 5A. The highly interacting part is highlighted in black box. (C) Alphafold3 interaction of the black box region from figure 5B, showing Jhc1 (magenta) and Ccdc33 c-terminal coiled-coil region (green). (D) PAE plot of figure 5C. The highly interacting part is highlighted in black box. (E) Jhc1 (magenta) and Ccdc33 c-terminal coiled-coil region (green) highlighted in figure 5D. ILE69 is highlighted in red. (F and G) Image of *Xenopus* MCC expressed with membrane-RFP (magenta) and GFP-Jhc1 (F)(green) or GFP-Jhc1-I69K (G)(green). Scale bar represents 10μm. (H) Quantification of GFP-Jhc1 (black) and GFP-Jhc1-I69K (magenta) fluorescent intensity along the axoneme. Graph shows mean ± SD (n=120 cilia, 40 cells, 10 embryos, 3 experiments).

We mutated Jhc1 Isoleucine-69 to Lysine, thereby mimicking Thtpa at this residue. The mutation in Jhc1 didn’t disrupt the stability of protein expression, which was confirmed with western blot analysis (Supp. Fig. 9). Excitingly, this point mutant completely failed to localize to the EDT of *Xenopus* cilia (Fig. 7F-H). These results are consistent with the possibility that Jhc1 is a pseudoenzyme derived from Thtpa, and the loss of the catalytic site in Thtpa facilitated novel protein-protein interaction and EDT localization in MCC cilia.

## Discussion

Here, we show that the previously uncharacterized protein Jhc1 plays a key organizing role at the EDT in *Xenopus* multi-ciliated cells (MCCs). Jhc1 interacts with Ccdc33 and is required to organize the distal region of cilia. Structural analysis and sequence alignment revealed that Jhc1 shares conserved secondary structure with Thtpa, a well-studied enzyme found in all eukaryotic lineages^36,38^, though Jhc1 is absent in mammals and non-vertebrate animals. Our data suggest the possibility that Jhc1 evolved from Thtpa where conserved catalytic residues were lost, and more interestingly, one of these residues became responsible for the localization of Jhc1 to the EDT.

Phylogenetic analysis across diverse species revealed Jhc1 as an independent clade in non-mammalian vertebrates (Supp. Fig. 2). This pattern raises the possibility that similar structural features evolved convergently or were maintained by similar functional constraints, despite the very divergent functions of Jhc1 and Thtpa. However, the deep sequence alignment suggests that Thtpa duplication and neofunctionalization gave rise to the pseudoenzyme Jhc1 which was later lost in mammals. This idea is supported by Jhc1 displaying loss of key catalytic residues present in Thtpa, yet retaining a similar secondary structure (Fig. 2B and Fig.6). Together, these findings indicate that diversity of the distal ciliary region may arise through functional divergence of conserved protein architectures, rather than the emergence of entirely novel structures.

The absence of Jhc1 in mammals may provide clues to distinct structures in distal ciliary regions across different species. For example, the ciliary tips of *Xenopus* embryonic epithelial MCCs possess a ball-like structure capping the central pair^20^, whereas the mammalian tracheal MCC cilia tip exhibits a flat plate-like structure linked to central pair and microtubule doublets together^11,17,47^. Both *Xenopus* and mouse motile cilia express tip proteins Ccdc78 and Ccdc33^20^, but show different morphologies^16^, potentially due to the presence of different components contributing to the tip complex, including Jhc1. Our findings thus provide new insights into evolution and the diversity of the tips of motile cilia.

## Methods

### Animal husbandry

All *Xenopus* experiments were conducted in accordance with the animal protocol AUP-2024-00130 and the animal ethics guidelines of the University of Texas at Austin. To induce the ovulation, female adult *Xenopus laevis* was injected with 400unit of hCG (Human chorionic gonadotropin) and incubated in a 16°C incubator for overnight. *In vitro* fertilization was performed by mixing the *Xenopus egg* with homogenized testis in 1/3X MMR (Marc’s Modified Ringer’s).

### Plasmid construction, mRNA synthesis and morpholino

Gene sequence was obtained from Xenbase (https://www.xenbase.org/) for *Xenopus laevis*, and NCBI (https://www.ncbi.nlm.nih.gov/) for *Danio rerio* and *Alligator sinensis*. Total RNA was extracted from *Xenopus laevis* embryo using TRIzol reagent (Invitrogen) and then reverse transcribed into cDNA using M-MLV Reverse Transcriptase (Invitrogen). cDNA of *Danio rerio* was a kind gift from Donghwa Suh in Gross lab at UT Austin. For Alligator coding sequence, gene block was synthesized from GenScript (https://www.genscript.com/). Coding sequences of genes were amplified by PCR using Q5® High-Fidelity DNA Polymerase (NEB). Amplified sequences and pCS10R vector containing fluorescence tag were digested with restriction enzymes and ligated using T4 DNA ligase. Ligated products were inserted into competent cells by transformation. Cloned constructs were linearized to synthesize mRNA using mMESSAGE mMACHINE™ SP6 Transcription Kit (Invitrogen).

Anti-sense morpholino of Ccdc78 and Ccdc33 were designed to block RNA splicing, morpholino of Jhc1 was designed to block translation based on the sequence from Xenbase database. The morpholinos were manufactured by Gene Tools. Morpholino sequences are as follows:

Ccdc33.L MO: 5’-GGTCAGGTAGTCACAGTATAAGAA-3’

Jhc1 MO : 5’-ATATAATCGGAGCGATCTGTTG -3’

### Microinjection in *Xenopus* embryos

Fertilized *Xenopus* embryos were de-jellied with 3% L-cysteine in 1/3X MMR (pH.7.8), washed and manipulated in 1/3X MMR. Embryos were manipulated in 2% ficoll during the injection. During 4-cell stage, 2 cells of ventral-animal side were injected with mRNA. For the Halo-tagged mRNA, ligands (Promega, Janelia Fluor® 646 HaloTag® Ligand) were injected together. The concentration of the mRNA is as follows: membrane-BFP 70pg/embryo, Spef1-RFP 45pg/embryo, GFP-Jhc1 25pg/embryo, GFP-Ccdc33 70pg/embryo, GFP-Alligator-Jhc1 70pg/embryo, GFP-fish-Jhc1 70pg/embryo.

### Image acquisition and analysis

For multi-ciliated cells live-imaging, *Xenopus* embryos were mounted between the cover glass with a small amount of 1/3X MMR and imaged immediately.

Confocal images were acquired with Zeiss LSM700 laser scanning confocal microscope using a plan-apochromat 63X/1.4 NA oil objective lens (Zeiss). Bright-field images were captured on a Zeiss stereomicroscope.

Quantitative measurement of images was done using Fiji. The fluorescent intensity values of each measured pixel for each protein were normalized with the average intensity of the entire distal-most four microns of individual cilia. Graph generation and statistical analysis including error bars, mean ± SD and P values were performed using Prism 10 software.

### Whole mount in situ hybridization

Whole mount in situ hybridization was performed based on previously described method^49^, using digoxigenin-labeled RNA probes against *jhc1* and *thtpa*. The anti-sense probe was transcribed using T7 RNA polymerase (NEB)

### PSI BLast analysis

Two iterations of PSI Blast were used to search the Refseq_protein databases for two mammals: *Mus musculus; Homo sapiens*; two reptiles: *Alligator mississippiens; Notechis scutatus*; one anuran and one urodele amphibian: *Xenopus tropicalis; Ambystoma mexicanum*; one cartilaginous and one bony fish, *Leucoraja erinacea; Danio rerio*, one cnidarian, *Nematostella vectensis*, and one Brachiopod: *Lingula anatine*. One iteration of PSI Blast retrieved only nine proteins, but two iterations retrieved twenty-four proteins. A third iteration did not retrieve additional proteins.

### Multiple sequence alignment

Protein sequences for Jhc1 and Thtpa were identified using PSI BLAST, as outlined above. As out groups, for a tree, we used two CYTH domain proteins identified by HHpred(cite) in search of the *Drosophila* and *C. elegans* proteomes.. Sequences were manually inspected and cleaned to remove ambiguous characters, frameshifts, and duplicates before analysis. Multiple sequence alignment was performed using MAFFT version 7 ^50^ with the L-INS-i strategy, which employs iterative refinement based on local pairwise alignments and excels for datasets with moderate divergence and conserved motifs amid variable regions. Maximum-likelihood phylogenetic reconstruction followed using IQ-TREE2 ^51^, with ModelFinder automatically selecting the best-fit substitution model and incorporating FreeRate (+R) modeling for site-specific rate heterogeneity. Node support was assessed via 1000 ultrafast bootstrap replicates integrated into the final treefile, capturing both primary similarities and subtle evolutionary rate patterns. The resulting treefile was imported into R using the ape ^52^ and visualized using ggtree^53^.

### Alphafold3 structure prediction

The protein structure predictions were predicted by Alphafold3 using Alphafold server (https://alphafoldserver.com/).

## Notes

### Competing Interest Statement

The authors have declared no competing interest.

### Funding

- NHLBI

