## Supplemental figures for "A pseudoenzyme controls the architecture of the extreme distal tip of motile cilia"

### Supplemental figure 1. Synteny of Jhc1 homologues

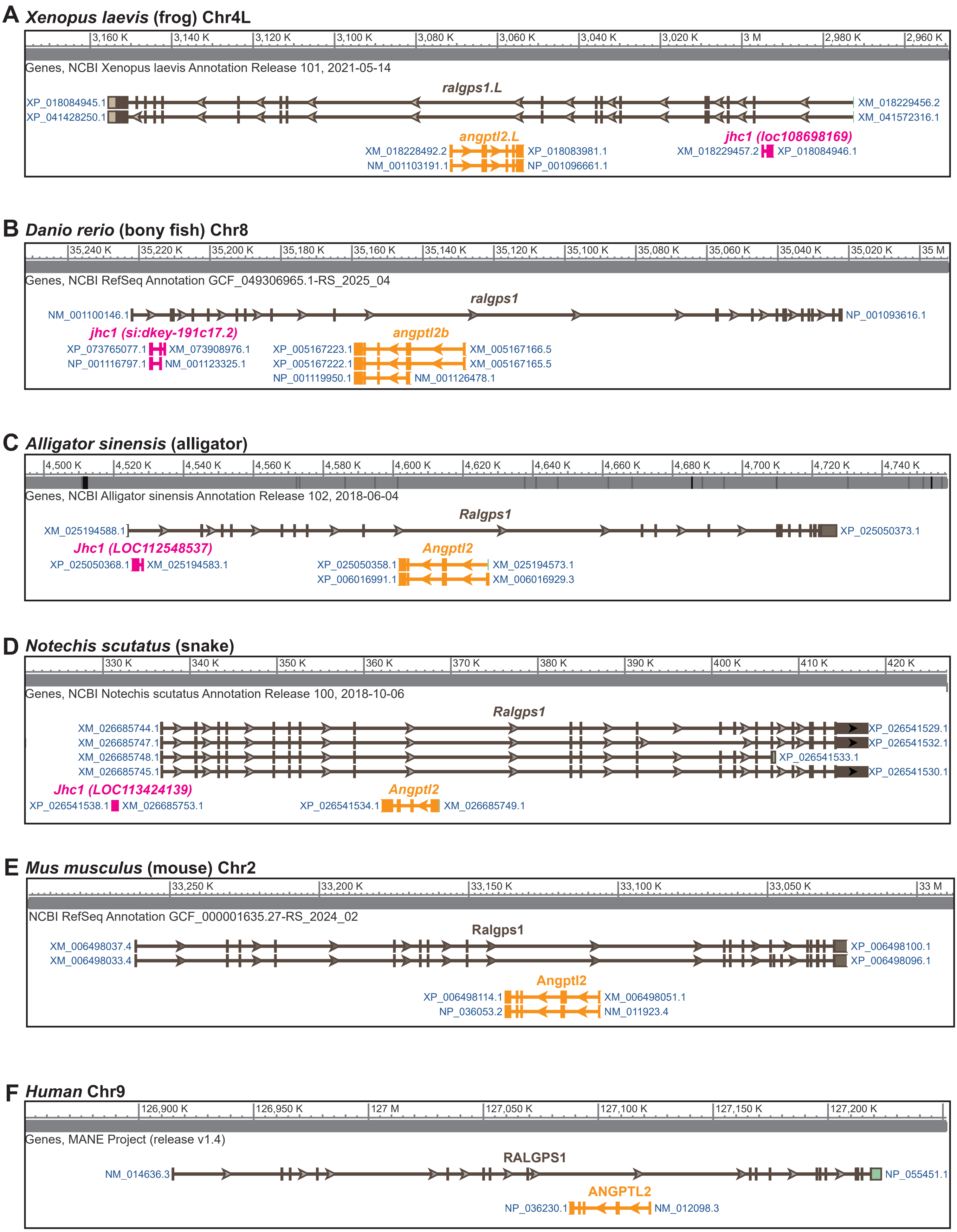

#### Supplemental figure 2. Multiple sequence alignment of Thtpa and Jhc1 across diverse species

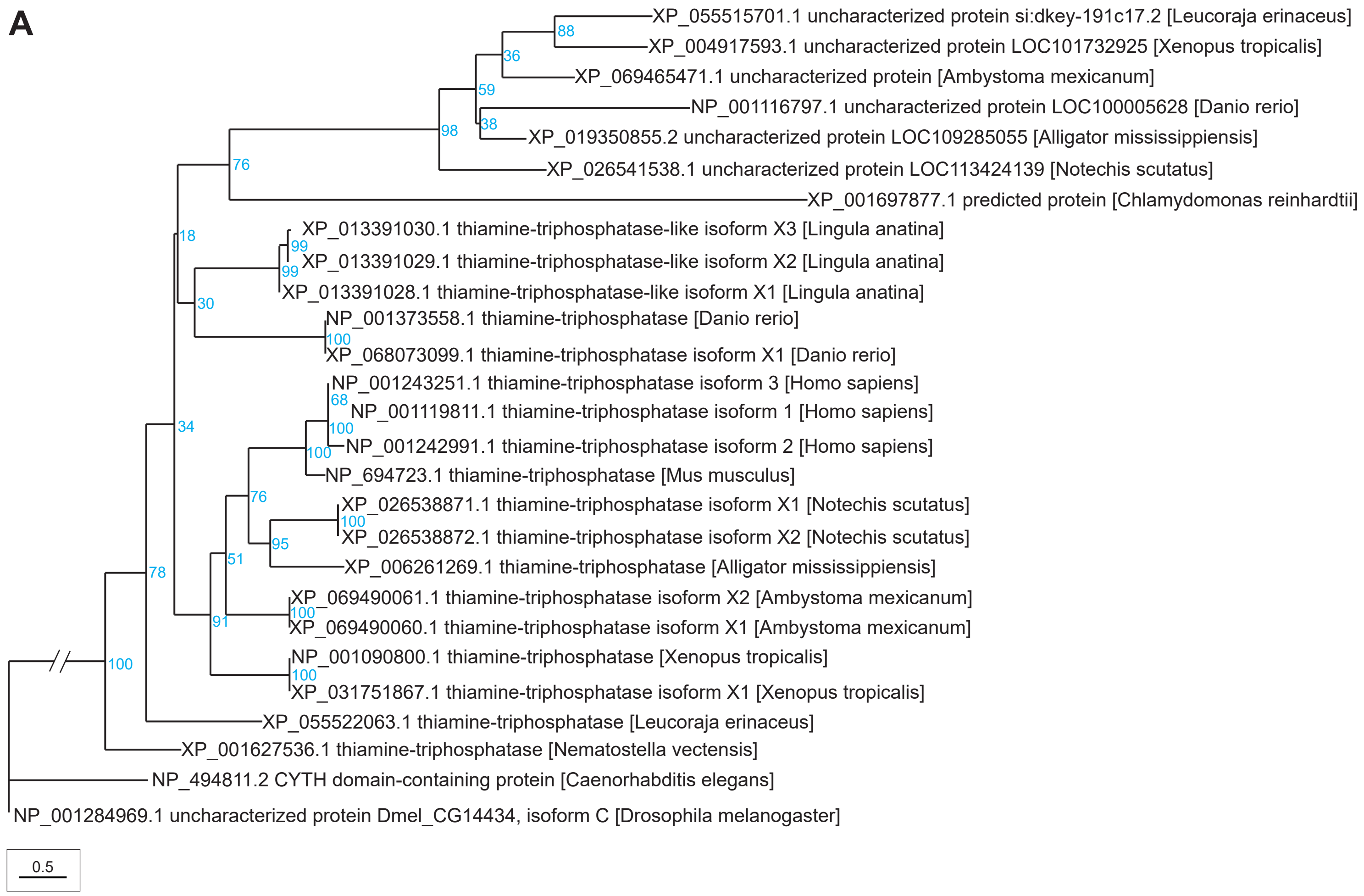

### Supplemental figure 3. Synteny of Thtpa homologues

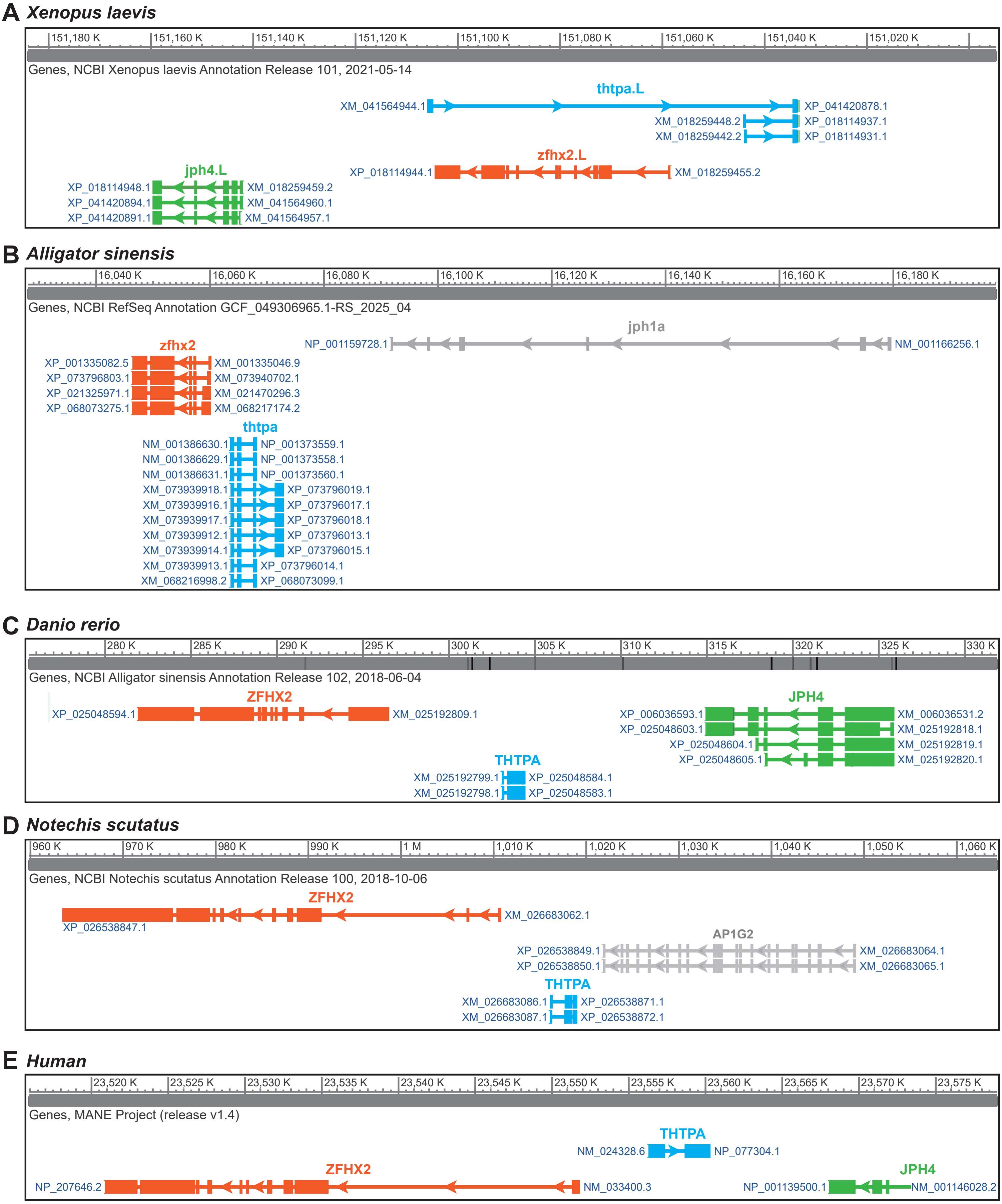

#### Supplemental figure 4. Comparison of secondary structures of Thtpa and Jhc1 across diverse species

A

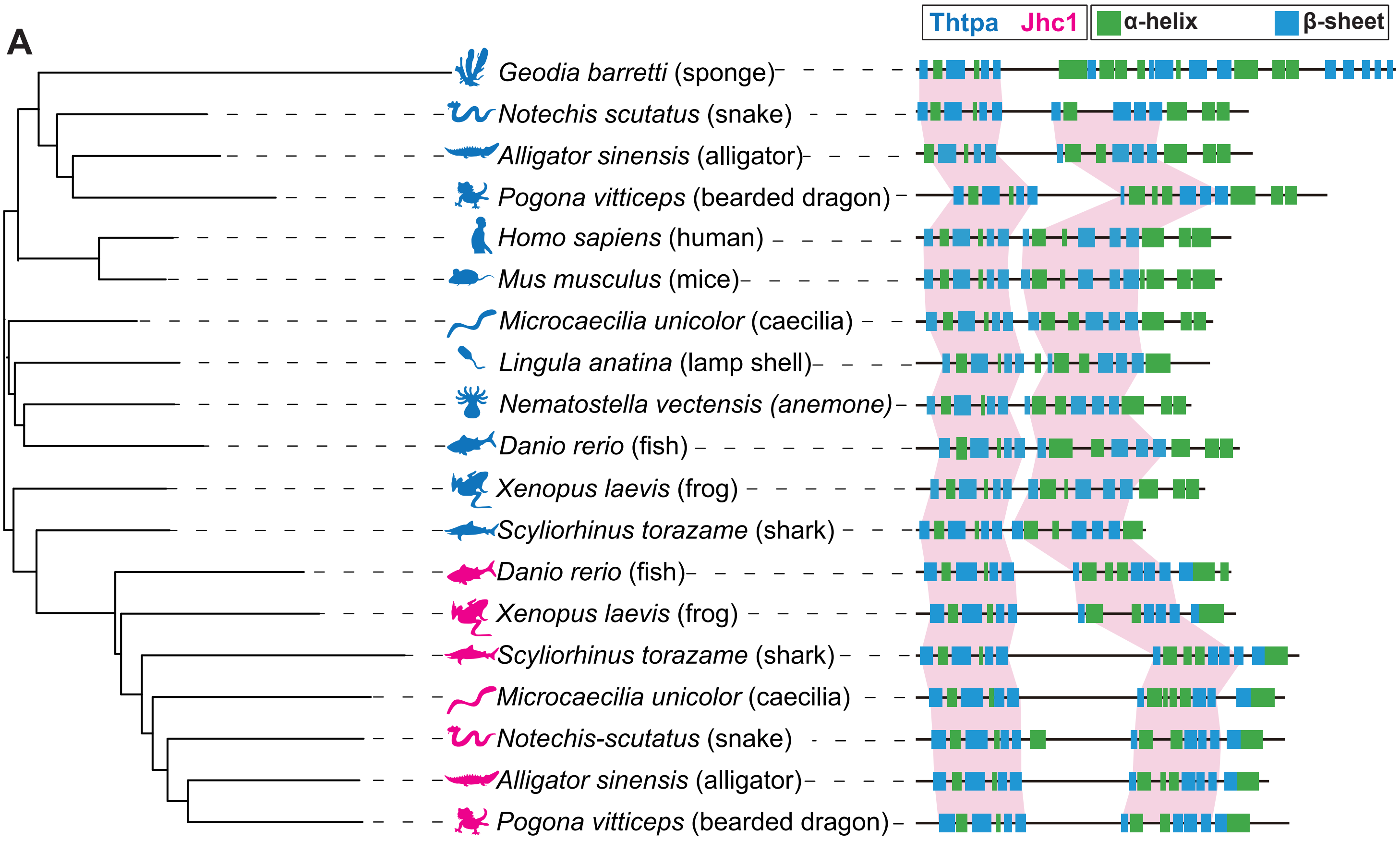

#### Supplemental figure 5. Structure of Thtpa from Alphafold3 prediction and PDB crystal structure

X-ray diffraction

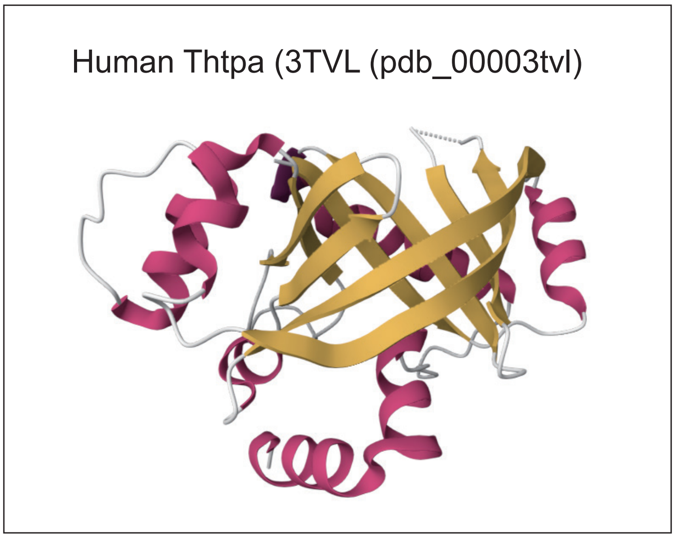


Alphafold3 prediction

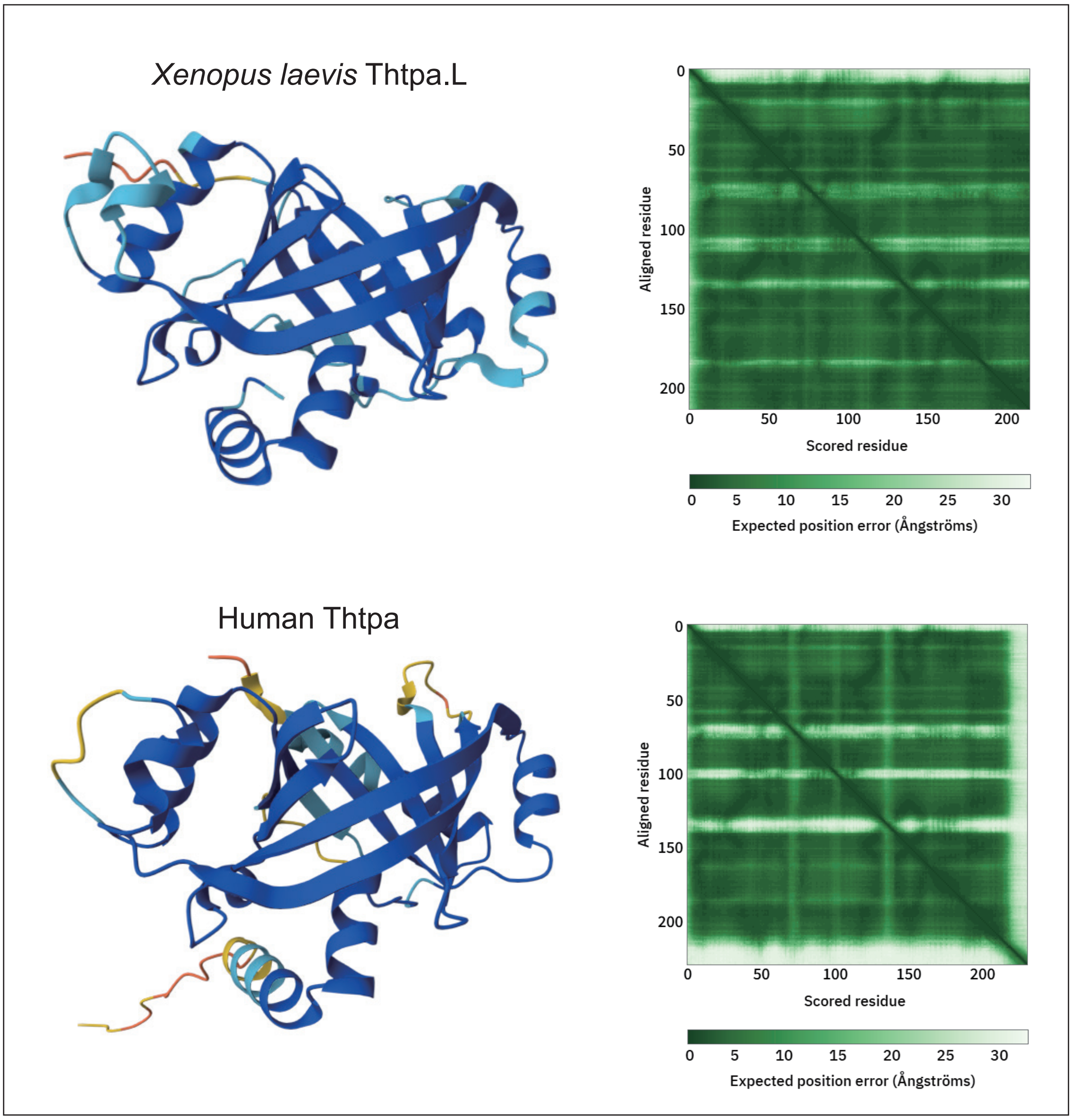

### Supplemental figure 6. Alphafold3 structure of fish and alligator Jhc1

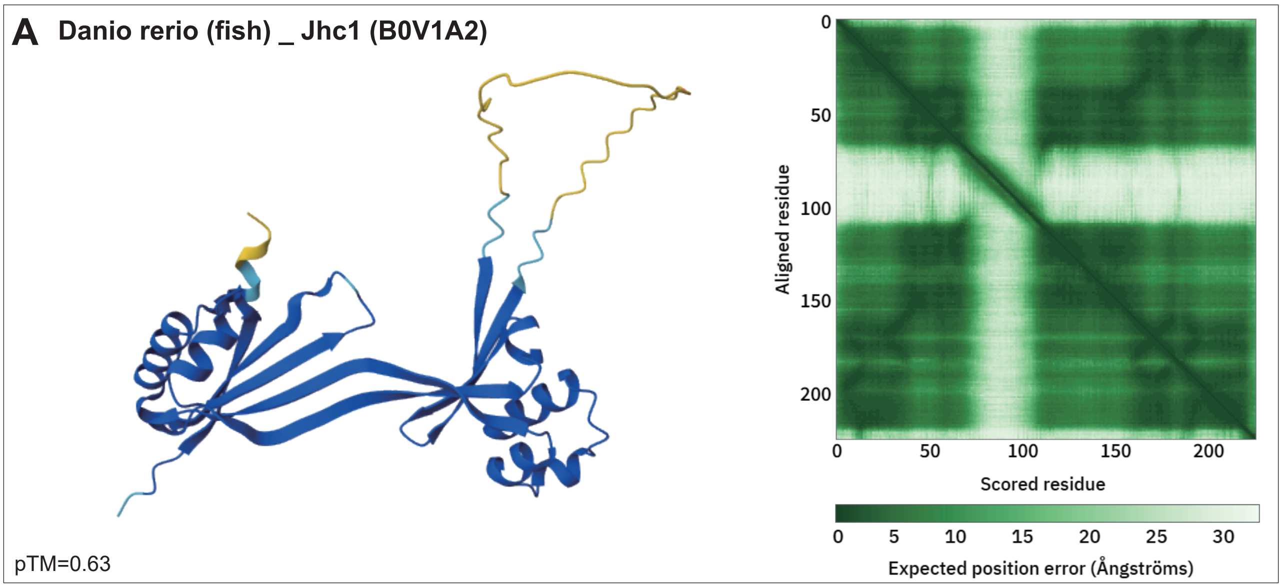


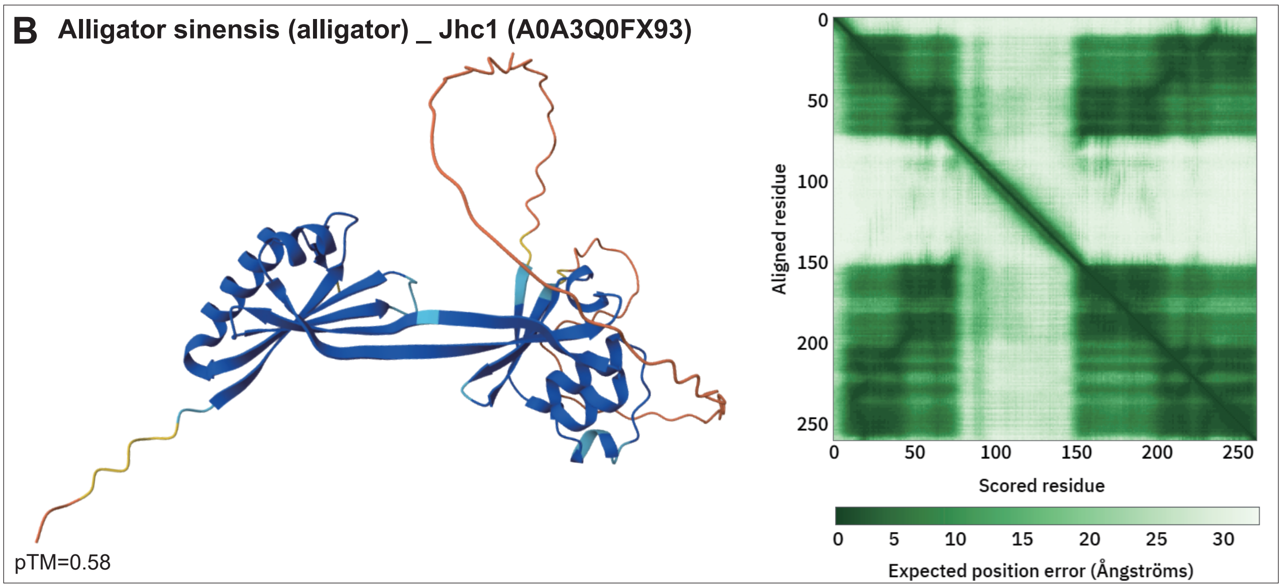

#### Supplemental figure 7. Localization of *Xenopus* and *Danio rerio* Thtpa in MCCs

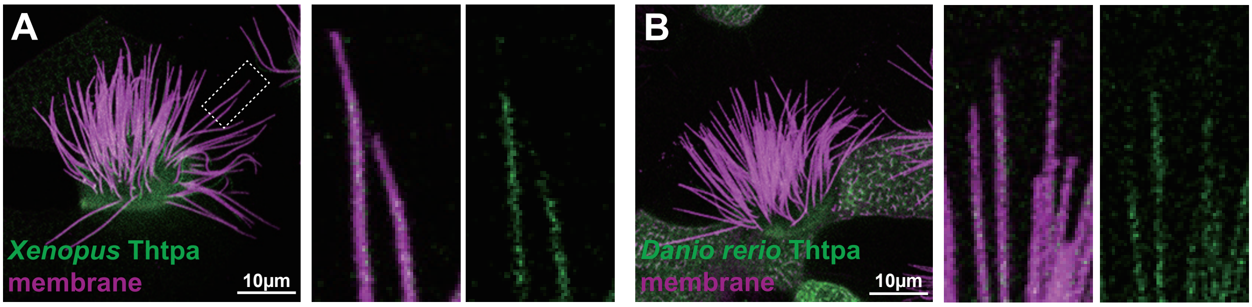

#### Supplemental figure 8. PAE plots of Mg2+ with Thtpa and Jhc1, catalytic residues in human and *Xenopus* Thtpa

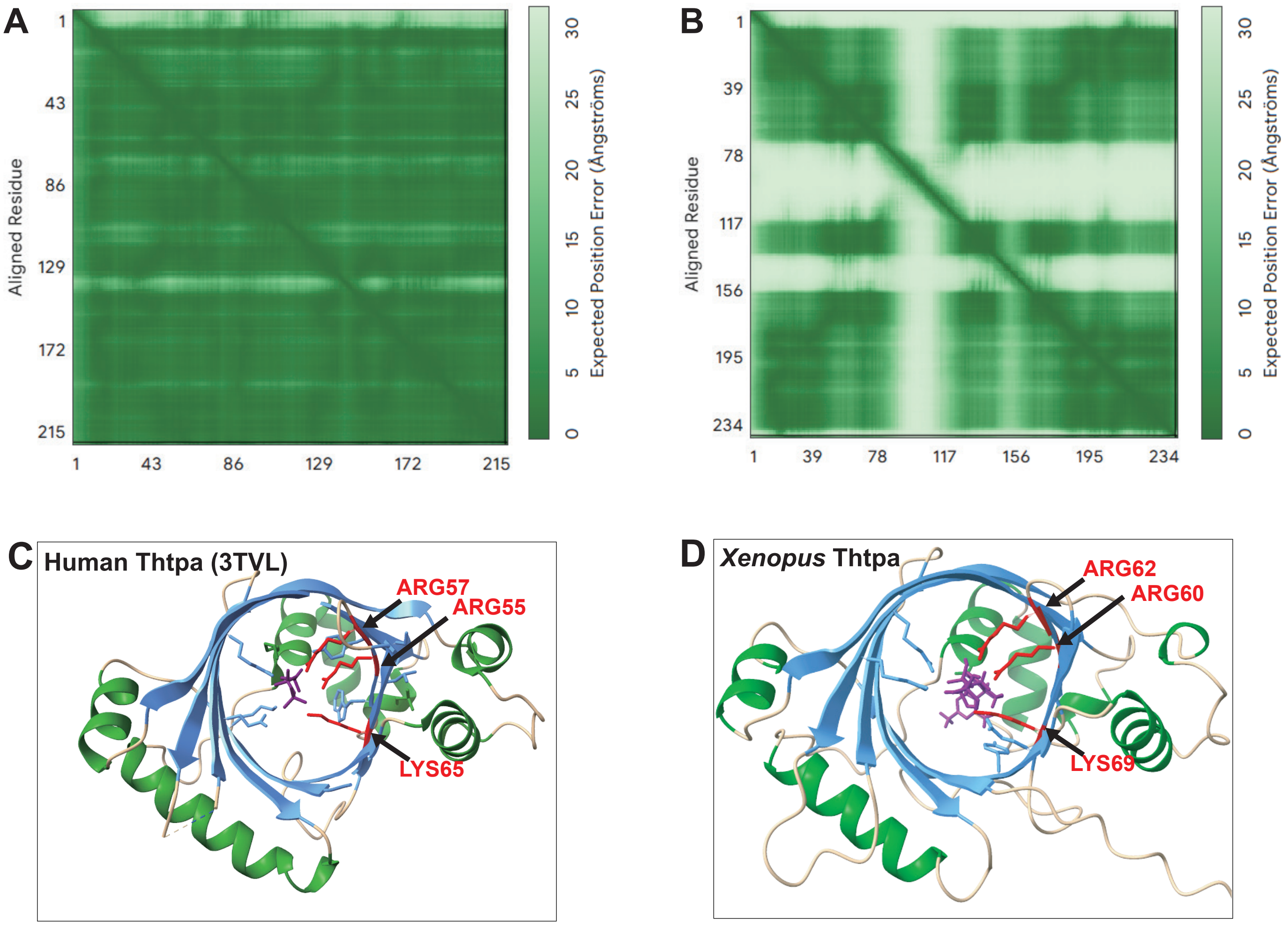

### Supplemental figure 9. Western blot of GFP-tagged Jhc1 and Jhc1-I69K

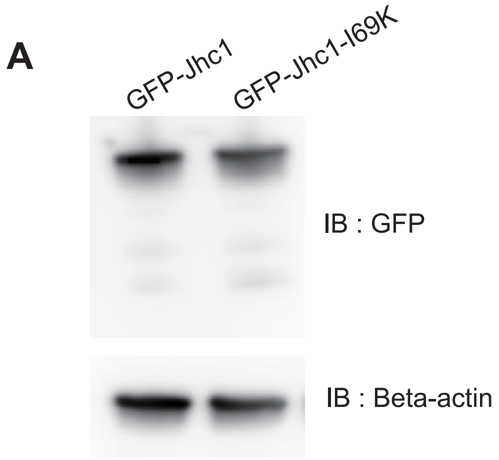

**Supplemental figure 1. Synteny of Jhc1 homologues**

(A-F) Genomic locus of *ralgps1* in *Xenopus laevis* (A), *Danio rerio* (B), *Alligator sinensis* (C), *Notechis scutatus* (D), mouse (E) and in human (F). The diagram shows the transcriptional orientations of *ralgps1* (brown) with neighboring genes *jhc1* (magenta) and *angpt2* (yellow) conserved across species. Human genome lacks *jhc1* gene.

**Supplemental figure 2. Multiple sequence alignment of Jhc1 and Thtpa across diverse species**

(A) Phylogenetic tree of multiple sequence alignment of Jhc1, Thtpa and CYTH-domain containing protein across diverse species. Bootstrap values are indicated at the branching points (1000 replicates). The scale bar represents 0.5 substitutions per site.

**Supplemental figure 3. Synteny of Thtpa homologues**

(A-E) Genomic locus of *thtpa* in *Xenopus laevis* (A), *Alligator sinensis* (B), *Danio rerio* (C), *Notechis scutatus* (D) and in human (E). The diagram shows the transcriptional orientations of *thtpa* (blue) with neighboring genes *zfhx2* (orange) and *jph4* (jph1a in *Alligator sinensis*)(green) conserved across species. Snake lacks jph4.

**Supplemental figure 4. Comparison of secondary structures of Jhc1 and Thtpa across diverse species**

(A) Secondary structure alignment of Jhc1 and Thtpa across diverse species. α-helix is shown in green and β-sheet in blue. Conserved structural regions, based on shared α-helix and β-sheet patterns, are highlighted in magenta.

**Supplemental figure 5. Structure of Thtpa from Alphafold3 prediction and PDB crystal structure**

Image of human Thtpa structure from PDB_0003tvl (Top box). Image of Alphafold3 prediction of human and *Xenopus* Thtpa with PAE plots (Bottom box). The crystal structure of Thtpa reflects the structure predicted from Alphafold3.

**Supplemental figure 6. Alphafold3 structure of fish and alligator Jhc1**

(A-B) Alphafold3-predicted structure of fish (B0V1A2)(A) and alligator (A0A3Q0FX93)(B) Jhc1 with corresponding PAE plots on the right.

**Supplemental figure 7. Localization of *Xenopus laevis* and *Danio rerio* Thtpa in *Xenopus* MCC**

(A and B) *Xenopus* MCC expressed with membrane-RFP (magenta) and *Xenopus* Thtpa (green)(A) or *Danio rerio* Thtpa (green)(B). The magnified view of the cilium is shown on right. Scale bar represents 10μm. Double slashes indicate truncated branch lengths.

**Supplemental figure 8. Alphafold3 prediction of Mg2+ with Thtpa and Jhc1**

(A-B) Alphafold3 prediction PAE plots of Thtpa (A) or Jhc1 (B) with Mg2+.

(C) Crystal structure of human Thtpa interacting with triphosphate (PDB_0003tvl), with conserved residues highlighted in red (ARG 55, ARG 57, LYS 65).

(D) ChimeraX prediction of *Xenopus* Thtpa interacting with triphosphate, with conserved residues highlighted in red (ARG 60, ARG 62, LYS 69).

**Supplemental figure 9. Western blot of GFP-tagged Jhc1 and Jhc1-I69K**

Western blots of GFP-Jhc1 and GFP-Jhc1-I69K from *Xenopus* lysates blotted with anti-GFP and anti-beta-actin.

**Supplementary table 1**. BLASTp analysis of Jhc1

**Supplementary table 2**. PSI-BLAST analysis of Jhc1

**Supplementary Table 3:** HHPred analysis of Jhc1

**Supplementary table 4**. BUDE alanine scan analysis result of Jhc1 and Ccdc33 coiled-coil region
